# *In vitro* activity of beauvericin against all developmental stages of *Sarcoptes scabiei*

**DOI:** 10.1101/814707

**Authors:** Charbel Al Khoury, Nabil Nemer, Georges Nemer, Mazen Kurban, Charlotte Bernigaud, Katja Fischer, Jacques Guillot

**Affiliations:** Faculty of Agricultural and Food Sciences, Holy Spirit University of Kaslik, P.O. Box 446 Jounieh, Lebanon; Research group Dynamyc, EA7380, Université Paris-Est Créteil, Ecole nationale vétérinaire d’Alfort, USC ANSES, 7 Avenue du Général de Gaulle, 94700 Maisons-Alfort, France; Department of Biochemistry and Molecular Genetics, American University of Beirut, Riad El Solh, Beirut, Lebanon; Department of Dermatology, American University of Beirut, Beirut, Lebanon; Department of Dermatology, Faculté de Médecine Paris-Est Créteil, AP-HP, Henri Mondor hospital, 51 Avenue du Maréchal de Lattre de Tassigny, 94010 Créteil, France; Scabies Laboratory, QIMR Berghofer Medical Research Institute, Infectious Diseases Program, Brisbane, Australia

**Keywords:** *Sarcoptes scabiei*, scabies, beauvericin, mycotoxin, treatment

## Abstract

**Background:** Scabies is a frequent cutaneous infection caused by the mite *Sarcoptes scabiei* in a large number of mammals including humans. As the resistance of *S. scabiei* against several chemical acaricides has been previously documented, the establishment of alternative and effective control molecules is required.

**Objectives:** In this study, the potential acaricidal activity of beauvericin was assessed against different life stages of *S. scabiei* var. *suis* and, in comparison with dimpylate and ivermectin, two commercially available molecules used for the treatment of *S. scabiei* infection in animals and/or humans.

**Methods:** In our *in vitro* model, developmental stages of *S. scabiei* have been placed in Petri dishes filled with Columbia agar supplemented with pig serum and different concentrations of the drugs. Moreover, the toxicity of beauvericin against cultured human fibroblast skin cells was evaluated using an MTT proliferation assay

**Results:** Beauvericin showed higher activity against adults and eggs of *S. scabiei* when compared to dimpylate and ivermectin. In addition, cell sensitivity assays demonstrated low toxicity of beauvericin against primary human fibroblast skin cells.

**Conclusion:** These results revealed that the use of beauvericin is promising and might be considered for the treatment of *S. scabiei* infection.

## Introduction

Scabies is a frequent cutaneous infection caused by the mite *Sarcoptes scabiei* in a large number of mammals including human.^1^ Human scabies was recently recognized as a neglected tropical disease by the WHO, due to its high global prevalence estimated to be around 100-200 million cases a year, ^2^ and high morbidity.^3^ Although primary infection with *S. scabiei* is limited to severe itching and allergic rash, secondary infections with bacteria such as group A streptococcus or *Staphylococcus aureus* could lead to severe acute infectious complications and even death,^4^ hence the importance of early and efficient eradication of the mites.^5^ There is a limited number of drugs that could be used for the treatment of scabies. Furthermore, the resistance to some of the drugs of *S. scabiei* is emerging, caused by re-infection or incorrect use of acaricides, may lead to an excessive and random use of treatments posing threat to patient health.^6^ Less susceptible populations of *S. scabiei* may emerge from the repeated exposure to a single type of acaricide. The resistance development may involve a mutation to the target site of the acaricide molecule or an up-regulation for genes encoding for detoxification enzymes (Van Leewan *et al.*, 2010). Therefore, the development of new acaricides with new mode of action against *S. scabiei* is required.

Many secondary metabolites produced by fungi have been used in medicine and agriculture.^7^ The entomopathogenic fungus *Beauveria bassiana* is known to produce beauvericin, a secondary metabolite belonging to the enniatin antibiotic family.^8^ This cyclic hexadepsipeptide was proven to have many biological effects including insecticidal, antitumor, antibacterial, and antifungal activity.^9^ Its mechanism of action is thought to be ionophore-induced apoptosis and DNA fragmentation.^7^ Recently, there is an ongoing interest for cyclic depsipeptide as topically applied medicines, treating per example, psoriasis, eczema and skin cancer (Cruz *et al.*, 2009). Accordingly, beauvericin could be considered as a potential new acaricide for the treatment of human and animal scabies. The objective of this study was to evaluate the *in vitro* efficacy of beauvericin against the different life stages of *S. scabiei*. In addition, this study evaluated the toxicity of beauvericin against cultured human fibroblast skin cells.

## Materials and method

### Ethics

All animals were maintained in strict accordance with good animal practices as defined by the French and European code of practice for the care and use of animals for scientific purposes (approval No. 02515.01). The biopsies were obtained after a written consent form was secured from each individual according to an approved protocol by the Institution Review Board (IRB) at the American University of Beirut (Protocol Number: DER.MK.01). The experiments were conducted in accordance with Good Clinical Practice and the ethical principles of the Helsinki Declaration

### *Sarcoptes* mites

*Sarcoptes scabiei* mites were collected from pigs maintained at CRBM (Centre de Recherche Bio Médicale), Maisons-Alfort, France. Pigs were experimentally-infected as described by Mounsey.^10^ Inoculation was done by directly introducing mite-infected skin crusts deep into the ear canals of five-week-old female piglets. Glucocorticoid treatment was initiated in naive piglets one week prior to inoculation and continued. For the present study, mites were collected from the pigs in weeks 15 and 16. Crusts in the external ear canal were gently removed and collected in a sterile Petri dish in the morning of the *in vitro* experiments. Mites crawled out of the crusts in about half an hour. Then they were picked one by one with a needle and under a dissecting stereomicroscope (Nikon©, SMZ645, Lisses, France).

### Molecules to be evaluated

Beauvericin 97% was purchased from Sigma-Aldrich. Dimpylate (diazinon) was purchased from Huvepharma™ (Segre en Anjou, France) (Dimpygale®, solution 100 mg/ml). Ivermectin was purchased from Boehringer-Ingelheim™ (Lyon, France) (Ivomec®, injectable solution 10mg/ml).

### In vitro efficacy tests on mites and eggs

To assess the efficacy of drugs (including beauvericin) against *S. scabiei* motile stages (larvae/nymphs and females), Petri dishes filled with Columbia agar supplemented with pig serum have been used for bioassays. To prepare the medium, 42g of Columbia agar (Bio-Rad, Marne-la-Coquette, France) were dissolved in 1L of distilled water. The solution was autoclaved for 15 min at 121°C then cooled down in a water bath at 53°C. Blood samples were obtained from pigs maintained in CRBM. Tubes of blood were centrifuged at 4500 rpm for 10 min at 4°C. The resulting supernatant was designated serum. For the preparation of each Petri dish, one ml of serum was added to 18 ml of Columbia agar medium at 53°C. Drugs to be tested were incorporated into the medium following the method described by Brimer^11,12^ with slight modifications. The mycotoxin beauvericin and two acaricide drugs (dimpylate and ivermectin) were tested with different concentrations. The absence of drug concentration was considered as negative control. The required volumes of serum-supplemented Columbia agar and molecules were pipetted in a tube and quickly transferred to a 9 cm sterile plastic Petri dish under a flow cabinet. Petri dishes were kept under the flow cabinet until the agar is solidified, and stored upside down at 4°C until use. The efficacy of 3 different concentrations 0.5, 5, and 50 μM was evaluated. Five females and five nymphs or larvae were inoculated in the middle of the plates and examined at 1, 2, 3, 4, 5, 6, 7, 8 and 24h after inoculation for survival assessment of motile stages at room temperature. Mites were considered dead when no movement occurred under the microscope during 5 min even after a gentle stimulation with a dissecting needle. After each inspection, mites were moved again to the center of the plate using a dissecting needle to lower chances of runaways.

To assess the efficacy of chemical products against *S. scabiei* eggs, 10 eggs were manually isolated using a fine needle and placed in the middle of beauvericin, dimpylate, or ivermectin-supplemented agar plates under the dissecting stereomicroscope as described above. Petri dishes were maintained at 37°C in an incubator for 5 days to promote egg development. Newly hatched larvae were recorded and removed from the Petri dishes.

Five replications were performed in three biological replicates, making it a total of 150 motile stages and 150 eggs observed for each treatment.

### Primary fibroblast culture

Human skin biopsies from healthy volunteer patients were delivered to the laboratory in culture medium (RPMI 1640: 450 ml, FBS: 50 ml, and penicillin-streptomycin solution x100: 1 ml). Each biopsy was transferred into a sterile Petri dish and rinsed with PBS to eliminate blood and debris. Two ml collagenase (Worthington™) were added to the medium before mincing the tissue with a scalpel. After incubation at 37°C for 1h, the digested tissue was transferred to a 15 ml conical tube and the Petri dish was rinsed twice with 2 ml of the medium and the liquid was collected in the same tube and span down at 200g for 5 min at room temperature. The pellet was washed twice with 3 ml of the medium to remove the collagenase, resuspended with 5 ml of the same medium and transferred to T25 flask. Finally, the cells were cultured in a 37°C humidified air incubator with 5% CO_2_. When there were sufficient cells, the latter were detached with trypsin and plated in another dish for further proliferation.

### Beauvericin cytotoxicity assessment

The cell death rates of treated fibroblast were used as an indicator to assess the cytotoxicity of beauvericin. Cultured cells were transferred to 96 wells plate and treated with 12 different concentrations of beauvericin in triplicates (0-50 μM) when they reached 50-60% confluence. Cells were continuously exposed to the drugs for 48h and subsequently assessed for cell death. The viability of the cells was assessed based on their metabolic activity using the MTT proliferation assay,^13^ as follows: 24h pre-treatment, fibroblast cells were starved in 100μl FBS free media. After starvation, fibroblast cells were exposed to the drugs as described above. Four hours prior to the end of the treatment, 10μl MTT dye (100 mg Thiazolyl Blue Tetrazolium Bromide, Sigma Aldrich, 20 ml PBS) were added to the wells. In addition, 100 μl MTT stop solution (12mM HCl, 0.05% isobutanol, 10% SDS) was added to cells before incubation at 37°C overnight. After 24h, the absorbance was measured at 550 nm on an ELISA plate reader. All tests were carried out in triplicates of three biological replicates.

### Statistical analyses

Efficacy data of all treatments against *S. scabiei* were analyzed by Kaplan Meier survival curves using software Statistical Package for the Social Sciences (SPSS, version 25).^14^ The statistical differences between data obtained with each treatment and the control for each experiment were measured by Log-rank test expressed by Chi-2 results and P-values (degree of freedom (df) = 1). P-value of ≤ 0.05 was considered significant. The lethal concentration (LC_50_) and lethal time (LT_50_) necessary to kill half of the mite’s population in addition to the lethal concentration (LC_50_) necessary to kill hald of fibroblast cells and their standard error were calculated using the probit regression analysis in (SPSS). The median time of 50% hatching (HT_50_) of the eggs was assessed.

LC_50_ of all treatments were analyzed by the statistical comparison test of means (ANOVA) using SPSS. The Tukey test was used at the 5% threshold for the separation of means.

## Results

The three molecules, beauvericin, dimpylate, and ivermectin were highly efficient against motile stages of *S. scabiei* mites (Table 1 and Figure 1). The mortality rates in the control group were below 5% during the first 8h post-exposure. The survival and hatching curves of mites and eggs exposed to different drugs are presented in Figure 1. In all tests, significant differences were found between each molecule and the control except for the tests against *S. scabiei* eggs with 0.5 μM of dimpylate and ivermectin. The highest mortality rates of all developmental stages were recorded with a concentration of 50 μM of all drugs. The efficiency of each treatment decreased steadily with the decrease in the concentration of the molecules: the lowest mortality rates were recorded within the plates supplemented with 0.5 μM of all drugs. Overall, a differential effect between the molecules and the concentration being used was notable 1h post-exposure (Table 1).

**Table 1:** Lethal time (LT_50_) to kill *Sarcoptes scabiei* females, nymphs and larvae or eggs and statistical differences between data obtained with each drug (beauvericin, dimpylate, and ivermectin).

|  |  | Females |  |  | Nymphs & larvae |  |  | Eggs |  |  |
| --- | --- | --- | --- | --- | --- | --- | --- | --- | --- | --- |
| Drugs and concentrations | | $X^2$ | $P$ | LT<sub>50</sub> ± S.E. (h) | $X^2$ | $P$ | LT<sub>50</sub> ± S.E. (h) | $X^2$ | $P$ | HT<sub>50</sub> ± S.E. (h) |
| Beauvericin | 0.5 µM | 112 | <0.05 | 3.4±0.1 | 34.4 | <0.05 | 4.7±0.2 | 39 | <0.05 | 2.3±0.1 |
|  | 5 µM | 159.7 | <0.05 | 1.9±0 | 79 | <0.05 | 4.1±0.2 | 54.2 | <0.05 | 2.5±0.1 |
|  | 50 µM | 165.9 | <0.05 | 1.4±0 | 124 | <0.05 | 2.3±0.1 | 85.9 | <0.05 | 2.9±0.1 |
| Dimpylate | 0.5 µM | 5.9 | <0.05 | 5.6±0.17 | 39.7 | <0.05 | 4.5±0.2 | 1.3 | >0.05 | 1.5±0 |
|  | 5 µM | 154.3 | <0.05 | 3.2±0.1 | 172 | <0.05 | 2±0.1 | 12.1 | <0.05 | 1.8±0 |
|  | 50 µM | 167 | <0.05 | 1.1±0 | 167.3 | <0.05 | 1.1±0 | 38.8 | <0.05 | 2.3±0.1 |
| Ivermectin | 0.5 µM | 151.4 | <0.05 | 2.8±0.1 | 166.5 | <0.05 | 1.6±0 | 0.3 | >0.05 | 1.4±0 |
|  | 5 µM | 156.8 | <0.05 | 2.4±0.1 | 166.9 | <0.05 | 1.2±0 | 6.5 | <0.05 | 1.6±0 |
|  | 50 µM | 166.5 | <0.05 | 1.6±0.1 | 164.3 | <0.05 | 1±0 | 19.8 | <0.05 | 1.9±0.1 |

Note: $X^2$ : Chi-square value; LT<sub>50</sub>: Lethal Time to kill 50% of mites; S.E.: Standard error; h: Hour; P: probability value; µM: micromolar.

**Table 2:** Concentrations of different molecules required to kill 50% of *S. scabiei* mites (females, nymphs/larvae and eggs).

|  | LC<sub>50</sub> (μM) ± S.E. |  |  |  |  |  |  |  |  |  |  |  |  |  |  |  |  |
| --- | --- | --- | --- | --- | --- | --- | --- | --- | --- | --- | --- | --- | --- | --- | --- | --- | --- |
| Exposure | 1h |  | 2h |  | 3h |  | 4h |  | 5h |  | 6h |  | 7h |  | 8h |  | 5 days |
|  | Females | Nymphs & larvae | Females | Nymphs & larvae | Females | Nymphs & larvae | Females | Nymphs & larvae | Females | Nymphs & larvae | Females | Nymphs & larvae | Females | Nymphs & larvae | Females | Nymphs & larvae | Eggs |
| Beauvericin | 34.7±<br>3.9<sup>c</sup> | 48.1<br>±1.7<sup>b</sup> | 7.8±<br>5.1<sup>a</sup> | 39.5±<br>3<sup>b</sup> | 2±0.9<br><sub>a</sub> | 28.4±<br>2.6<sup>b</sup> | 0.7±0<br>.2<sup>b</sup> | 18.9±<br>1.9<sup>b</sup> | 0.3±0<br><sub>b</sub> | 13.7±<br>1.3<sup>b</sup> | 0.3±0<br><sub>b</sub> | 10.4±<br>1.2<sup>b</sup> | 0.3±0<br><sub>a</sub> | 1.7±0.<br>6<sup>b</sup> | 0.3±0<sup>a</sup> | 0.9±0<sup>a</sup> | 77.5<br>±1.8<sup>a</sup> |
| Dimpylate | 7.09±<br>0<sup>a</sup> | 6.9±<br>1<sup>a</sup> | 5.9±<br>0.2<sup>a</sup> | 3.3±0<br>.2<sup>a</sup> | 4.1±0<br>.1<sup>a</sup> | 2.3±0<br>.7<sup>a</sup> | 3.9±0<br>.1<sup>a</sup> | 1.7±0<br><sub>a</sub> | 2.5±0<br>.2<sup>a</sup> | 0.5±0<br><sub>a</sub> | 1.5±0<br>.1<sup>a</sup> | 0.5±0<br><sub>a</sub> | 1.1±0<br>.4<sup>a</sup> | 0.4±0<br><sub>a</sub> | 0.8±0.<br>1<sup>b</sup> | 0.4±0<sup>a</sup> | 77.5<br>±1.8<sup>a</sup> |
| Ivermectin | 45.1±<br>0.9<sup>b</sup> | 5.7±<br>0.8<sup>a</sup> | 26±1<br>.5<sup>b</sup> | 0.4±0<br><sub>a</sub> | 4.7±3<br>.9<sup>a</sup> | 0.3±0<br><sub>a</sub> | 0.3±0<br><sub>b</sub> | 0.3±0<br><sub>a</sub> | 0.3±0<br><sub>b</sub> | 0.3±0<br><sub>a</sub> | 0.3±0<br><sub>b</sub> | 0.3±0<br><sub>a</sub> | 0.3±0<br><sub>a</sub> | 0.3±0<br><sub>a</sub> | 0.3±0<sup>a</sup> | 0.3±0<sup>a</sup> | 202.6<br>±16.5<sup>b</sup> |

Note: LC<sub>50</sub>: Lethal Concentration to kill 50% of mites; S.E.: Standard error; h: Hour; μM: micromolar \*Values followed by the same letter in the same column are not significantly different at the 5% threshold

**Figure 1:**
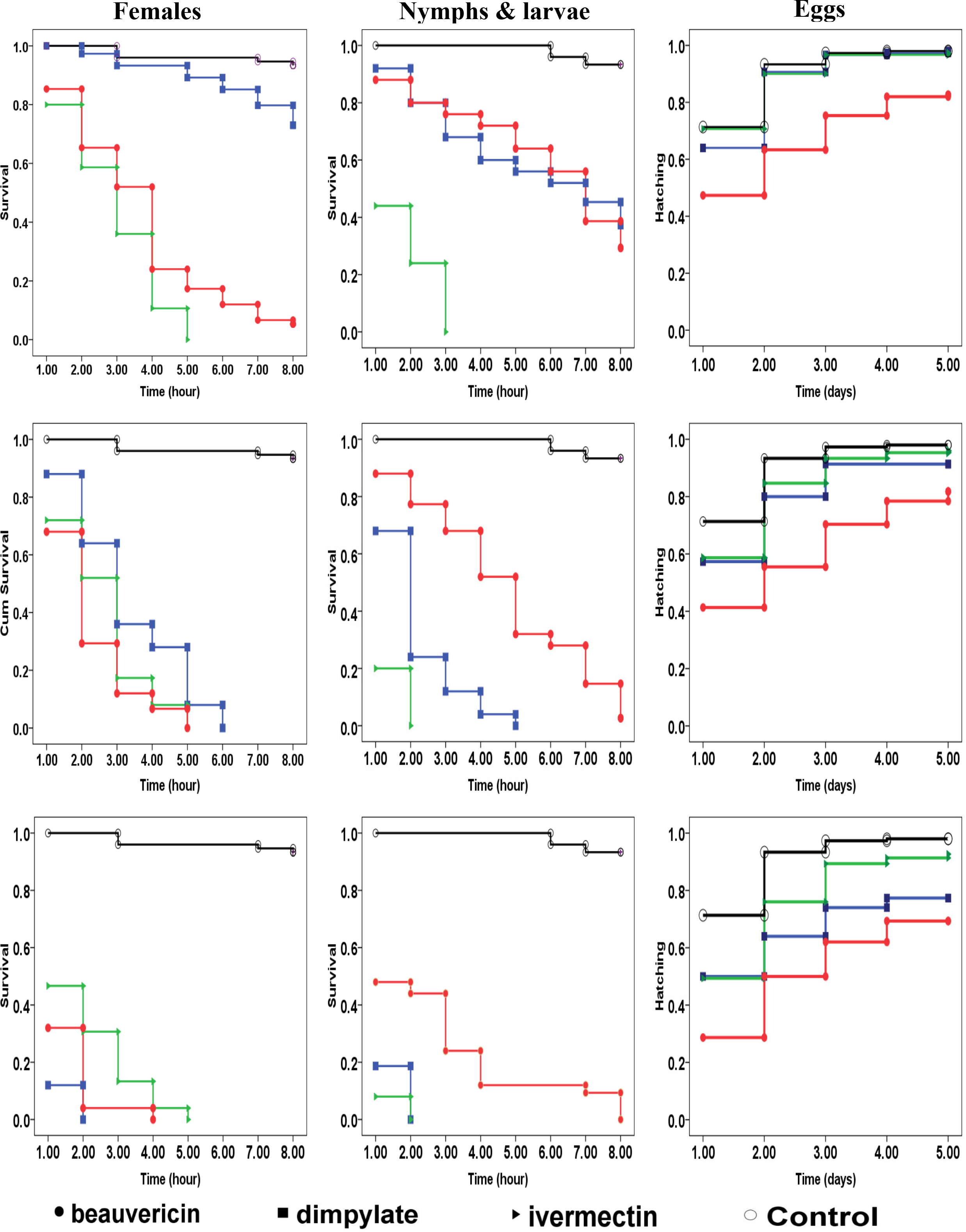
Curves representing survival (of motile stages) and hatching (of eggs) of *S. scabiei* exposed to acaricide molecules at different concentrations: a) 0.5 μM, b) 5 μM, and c) 50 μM. For each treatment, observed survivals and hatchings are presented using curves with markers; filled circle: beauvericin; filled square: dimpylate; filled triangle: ivermectin; open circle: control

The efficacy of the molecules at different concentrations can be put into the following order based on their chi-square value: beauvericin > ivermectin > dimpylate and ivermectin > dimpylate > beauvericin against females and immature forms, respectively (Table 1). The survival capacity seemed to be different according to the developmental stage of the mites. Dimpylate and ivermectin had higher efficacy on *S. scabiei* immature motile stages when compared to females; whereas, beauvericin displayed a higher efficacy on females at all concentrations (Table 1).

The activity of all three molecules against *S. scabiei* eggs was evaluated for 5 days. Results obtained with beauvericin, dimpylate or ivermectin were significantly different from those in the control group (Table 1). Among all the molecules tested against the eggs of *S. scabiei*, beauvericin demonstrated the best inhibition of hatching effect, when testing a concentration of 50 μM. The second-highest activity was recorded when treating eggs with the same concentration of dimpylate. The lowest activity was recorded at a concentration of 0.5 μM of dimpylate or ivermectin with no significant statistical difference compared to the negative control (Table 1).

LT_50_ values were different between treatment groups against the motile stages of *S. scabiei* (Table 1). The highest LT_50_ values (5.6 and 4.7h) were observed with a concentration of 0.5 μM of beauvericin and dimpylate against females and nymphs/larvae, respectively. The lowest LT_50_ values (1.1 and 1 h) were observed with a concentration of 50 μM of dimpylate and ivermectin against females and nymphs/larvae, respectively. The median time for hatching of 50% of the eggs was also recorded in this study (Table 1).

A significant difference was recorded between LC_50_ values of the drugs at 1h (F = 68.779, df = 2, P <0.05), 2h (F = 12.809, df = 2, P <0.05), 4h (F = 145.902, df = 2, P <0.05), 5h (F = 59.758, df = 2, P <0.05), and 6h- post exposure against females (F = 12.809, df = 2, P <0.05). The effect of the drugs was not notable 3h post exposure against females (F = 0.39, df = 2, P >0.05). A significant difference was also notable between LC_50_ values of molecules at any given time of the test against larvae: 1h (F = 367.273, df = 2, P <0.05), 2h (F = 1423.942, df = 2, P <0.05), 3h (F = 108.356, df = 2, P <0.05), 4h (F = 86.263, df = 2, P <0.05), 5h (F = 94.776, df = 2, P <0.05), and 6h post exposure (F = 68.387, df = 2, P <0.05). Median times of 50% hatching of the eggs are presented in Table 1. The highest median time for hatching (2.9 days) was recorded with a concentration of 50 μM of beauvericin. A significant difference was recorded between LC_50_ values of the molecules against eggs at 5 days post-exposure (F = 42.709, df = 2, P <0.05).

The cytotoxic effect of beauvericin was assessed at 48h post-exposure. The mycotoxin caused a dose dependent reduction in cell viability and lethal concentration 50 was calculated. The human fibroblast cells were moderately sensitive to beauvericin toxicity and LC_50_ was 4.8 μM.

## Discussion

The mortality rate in the control groups was low (< 5%) during the first 8h of the test indicating suitable conditions of the bio-assays. The survival of *S. scabiei* outside its host is considered to be the first limitation of *in vivo* studies. After 24h, the “natural” mortality rate at room temperature reached 19% which indicates that mortalities recorded at 24h may not be caused only by the acaricide effect of the tested molecules; therefore, results recorded at 24h post inoculation are discarded.

The efficacies of macrocyclic lactones (including ivermectin) or organophosphates (like dimpylate) against *S. scabiei* have been previously evaluated.^12,15,19,22^ However, most studies evaluated the efficacy of such molecules against motile stages of *S. scabiei* with no differentiation of the life stage. Studies evaluating the efficacy of acaricides against specific life stages of *S. scabiei* are few. The present study demonstrated that the application of dimpylate and ivermectin caused higher survival rates among females compared to immature forms. These results are in accordance with those from Mounsey^17^ who observed that *S. scabiei* nymphs and larvae were more vulnerable than females to ivermectin and moxidectin. Generally, larvae are more sensitive due to their greater surface area/volume ratio meaning more drug is absorbed through the cuticle during *in vitro* exposure. On the contrary, in the present study, *S. scabiei* females seemed to be more vulnerable to beauvericin than immature forms. These results are in accordance with those from Fu-Xing *et al.*, 2002 reporting a higher induction of two kinds of insecticides metabolic detoxifying enzymes by larvae of *Musca domestica* when compared to adults.

The resistance of *S. scabiei* to commercial products exists and might increase in the future; whereas, treatment failures of scabies infections in animals and humans have been previously reported.^20–23^ Beauvericin is known to be an ionophoric cyclodepsipeptide which forms complexes with cations and increases the permeability of biological membranes.^24–26^ Given the non-similarity of its mode of action to that of the commonly used neuro inhibitors, a cross-resistance of *S. scabiei* mites against beauvericin is unlikely to happen.

The three drugs selected for the bioassays had a low LT_50_ value, indicating a rapid effect on the mites. For beauvericin and dimpylate, LT_50_ values were related to the concentration of the molecules (death occurred more rapidly among mites treated with higher concentrations).

The dose-response test recorded low LC_50_ values indicating high efficacy of all drugs at killing *S. scabiei* mites. Moreover, LC_50_ values obtained in the present study were low when compared to those obtained by Mounsey.^17^ The latter study reported that 50.5 μM of ivermectin is required to kill 50% of *S. scabiei* mites 1h post-exposure (versus 45.1 μM in our study). This result could be explained by the fact that the strain of *S. scabiei* and/or the assay conditions used in the present study were different.

The next generation of scabicide molecules needs to target *S. scabiei* eggs and ensuing developmental stages. In the present study, high hatching rates were recorded among the eggs in contact with dimpylate and ivermectin. These results are in accordance with Dourmishev^27^, Usha and Nair^28^ demonstrating that presently used acaricides have a very limited inhibition of hatching activity. The present study demonstrated that beauvericin was moderately active against the eggs of *S. scabiei.*

Data about beauvericin cytotoxicity, especially against skin cell line are lacking. This lack of data could explain why there are no maximum guidance levels. The present study assessed for the first time the cytotoxic activity of beauvericin against primary fibroblast human skin cells. Heilos *et al.*^30^ investigated the cytotoxic activity of beauvericin against the principal constituent of the epidermis, HEK2 keratinocyte (IC_50_ = 5.4 μM). Furthermore, systemic kinetics and effect of beauvericin should also be assessed since the cyclic depsipeptide is transdermally absorbed^31^.

The impact of beauvericin was also assessed against several nucleated human cells (IC50s included human intestinal cell line Caco 2= 3.9μM; human liver cell line HEPG2 liver= 3.4μM; human normal vascular endothelial cells HUVEC = 2.4μM)^30^. The latter reviews revealed cytotoxic activity against human cell lines at relatively low concentrations; nevertheless, the present study demonstrated that the therapeutic index of beauvericin for scabies infection could be high. The susceptible dose against motile stages of *S. scabiei* are extremely low when compared that of all human cell lines. Furthermore, a study conducted by Taevernier *et al*.^31^ demonstrated that beauvericin concentration was 21 times higher in the epidermis than in the dermis after topical application of the mycotoxin. Moreover, transdermal kinetics is mediated by the outermost layer of the skin providing a protective reservoir for cyclic depsipeptides.^31^

In conclusion, this study presented the first evidence of *in vitro* efficacy of the mycotoxin beauvericin against different developmental stages of *S. scabiei* mites. Beauvericin showed higher efficacy against females and eggs of *S. scabiei* when compared to the two commercially available acaricides, dimpylate, and ivermectin. Furthermore, beauvericin had low cytotoxicity against fibroblasts. These preliminary results indicated that beauvericin may be considered as a new scabicide molecule. Further studies assessing the possibility of beauvericin application to treat scabies in humans or sarcoptic mange in animals are required. Bioavailability, toxicokinetic properties, distribution, absorption, metabolization and excretion of the mycotoxin need now to be documented. Studies measuring and confirming that the maximum concentration of beauvericin in the skin of patients are within the *in vitro* susceptibility range are crucial.

## Acknowledgment

We are grateful to Radia Guechi, and Francis Moreau for their laboratory assistance

## Funding

This work was funded by the Lebanese National Council for Scientific Research (CNRS), grant name (CNRS-L/USEK) and “Coopération pour l’évaluation et le développement de la recherche” CEDRE grant number 37349SA.

## Transparency declarations (conflicts of interest)

None to declare

